# A lectin that targets specific bacteria for killing on DNA-based extracellular traps

**DOI:** 10.1101/2020.05.04.077404

**Authors:** Timothy Farinholt, Christopher Dinh, Adam Kuspa

## Abstract

Animal defenses directed against bacteria include DNA-based extracellular traps (ETs) that are produced by innate immune cells. ET-bound bacteria are prevented from further tissue dissemination and are eventually killed by ET-bound antibacterial proteins. It is unclear how bacteria bind to ETs, though it has been proposed that the negatively-charged DNA scaffold of the ETs is involved. We have found that the bacterial-binding lectin CadA is a component of the ETs produced by the innate immune cells of *Dictyostelium discoideum* and is required for the binding and killing of two *Enterobacteriaceae* by ETs, but not other bacteria. Our results suggest that ETs selectively sequester bacteria and that lectins can facilitate bacterial killing by acting as ET-bacteria binding proteins.

## Introduction

The innate immune system consists of motile tissue-resident cells and freely circulating cells that move towards chemical signals released by cells at the site of acute infection (1, 2). They phagocytose microbes, secrete antimicrobial proteins and peptides, and release reticulated DNA-based ETs to eliminate pathogens. Neutrophils (3), eosinophils (4), and basophils (5) release antimicrobial proteins, such as myeloperoxidase and neutrophil elastase, either as cytoplasmic granules or bound to ETs derived from mitochondrial DNA (3,6–8). The reticulated DNA of ETs provides the structure for bacteria to bind to and to be exposed to a high local concentration of antimicrobials and this is presumably necessary for efficient bacterial killing. The social amoebae *D. discoideum* faces similar challenges from bacteria when it forms a multicellular organism in the soil (9). On it’s way to forming a spore forming fruiting body, aggregated amoebae form a motile slug that moves to the surface of the soil. This journey has the potential to expose the slug to pathogens that could reduce spore yields and overall fitness. To combat this, amoeba maintain a stable population of innate immune Sentinel cells (10). Sentinel cells retain the ability to phagocytize bacteria and they also release extracellular traps of a form and function similar to mammalian ETs (11). Amoebal ETs are released in response to bacteria and trap and kill bacteria, eventually being released behind the slug in the slime trail. *D. discoideum* ET release also requires reactive oxygen species produced by NADPH oxidase and signaling through *tirA*, a Toll/Interleukin-1 domain protein (11).

Studies have focused on identifying ET-associated antimicrobial proteins that decorate the structure (12), or elucidating the release pathway, but our understanding of how bacteria bind to ETs remains elusive. We describe the identification of a lectin, CadA, which is required for the binding of some bacterial species, but not others, to ETs. CadA is required for the efficient killing of those bacteria that require CadA for ET binding. Uncovering the mechanism(s) of bacterial binding to *D. discoideum* ETs may illuminate the specificity of amoebal ET function and this may improve our understanding of human ET function (7, 8).

## Materials and Methods

### Strains, growth, and plasmids

*Klebsiella pneumoniae* (*K.p*.) was grown in Standard Media (SM) or on SM agar (13). *Bacillus subtilis* GFP and *Pseudomonas aeruginosa* PAO1 were grown in LB media or on LB plates. *Dictyostelium discoideum* strains were derived from the axenic laboratory strain AX4 (14, 15), and maintained in HL5 media with 50 U/mL penicillin and 50 µg/mL streptomycin (Penn/Strep) or in co-culture with K.p. on SM agar plates [2% Bacto agar (BD difco) with SM media; 13]. Blasticidin-resistant *D. discoideum* strains (bs^r^) were grown in HL5 media supplemented with 4 µg/ml Blasticidin (ThermoFisher). The creation of the *cadA*-mutant strain was described previously (16).

### ET Production

Axenic *D. discoideum* cells were harvested at 1-2 × 10^6^ cells/ml. Cells were washed twice with Sorenson’s buffer (2.0g KH_2_PO_4_, 0.29g Na_2_HPO_4_ per liter, Sor) and resuspended to 1×10^7^ cells ml^-1^. To form slugs, 400 µl of cells were deposited onto a 10 cm plate of water agar (1.2% noble agar, with 2% ethidium bromide for Sentinel cell purification) and allowed to dry slightly. These plates were wrapped in tin foil and placed in a dark humid chamber to develop at room temperature (22°C). At 16 hours, slugs were harvested by scrapping into Sor buffer. Slugs were disaggregated then put through a 40-µm mesh cell strainer (Fisher Scientific) to remove clumps of cells. Samples are then aliquoted into 24-well plates or onto coverslips. Lipopolysaccharide purified from *K. pneumoniae* (LPS^*K.p*.^, 2-5 µg ml^-1^, Sigma Aldrich) or overnight bacterial cultures were added and incubated for one hour. ETs are then visualized by the DNA stain DAPI (Thermo Scientific). For biochemical analysis of ET-associated proteins, Sentinel cells were first purified using fluorescence activated cell sorting (FACS) and by selecting the brightest 1-2% of cells from the suspensions of slugs cells described above, as we have described previously (10). ETs were purified as described previously (11). Briefly, aliquots of purified Sentinel cells were incubated in standard micro-centrifuge tubes and exposed to buffer or LPS^*K.p*.^ (5 µg ml^-1^) for 3 hours to elicit ET production. The ETs readily stuck to the walls of the tubes and were separated from the Sentinel cells by low-speed centrifugation (5 min at 1,200*g*). The cell pellets were removed from the tubes and the ETs remaining on the walls of the tubes were solubilized in SDS-polyacrylamide gel electrophoresis (SDS-PAGE) sample buffer, and separated by SDS-PAGE (17).

### Biochemical methods

CadA was identified by the Mass Spectrometry Proteomics facility within the Advanced Technology Core laboratories of Baylor College of Medicine (www.bcm.edu/corelabs) using an excised coomassie stained band from an SDS-PAGE gel, similar to the one indicated in Fig. 1a. SDS-PAGE was carried out on 12% gels with BioRad 30% acrylamide/bis-acrylamide (29:1) solution. Protein bands were visualized by soaking gels in Coommassie R-250. To visualize CadA on ETs, ETs were developed on coverslips as above. Samples were incubated with anti-CadA monoclonal antibody MLJ11 for 30 minutes followed by three washes. MLJ11 was obtained from the laboratory of William F. Loomis (Univ. of Calif. San Diego). Secondary Alexa Fluor 488 conjugated goat anti-mouse antibodies (1:10,000) were then incubated for 30 minutes before three washes and imaging by fluorescence microscopy.

**Figure 1.**
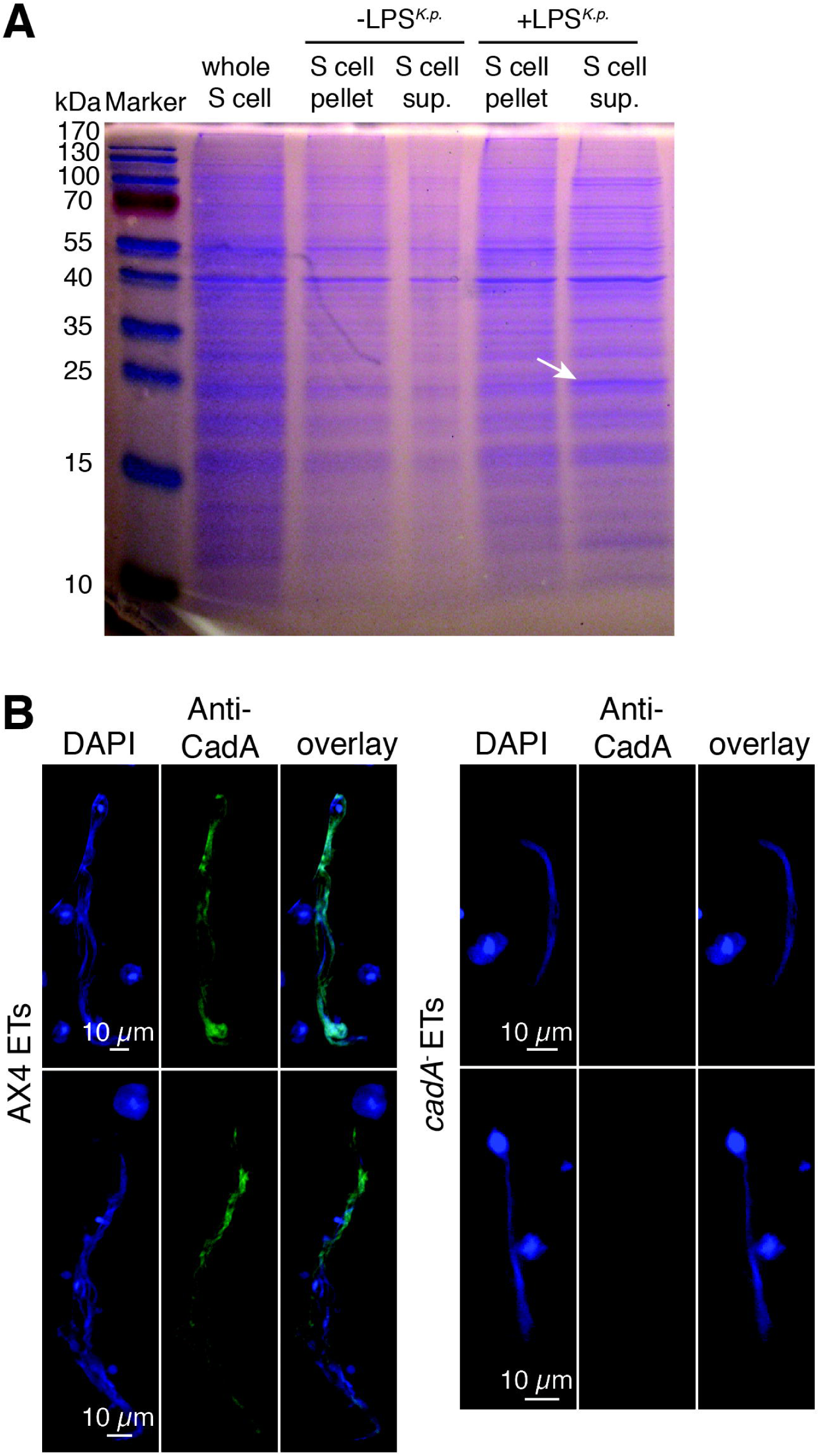
CadA lectin is associated with *D. discoideum* extracellular traps. (**A**) Sentinel cells were purified from disaggregated slugs and exposed to buffer with or without *K. pneumoniae* LPS (LPS^*K.p*.^) to elicit ET production. Samples were separated by centrifugation into a cell pellet fraction and a supernatant (sup.) fraction containing ETs, and the proteins were resolved by SDS polyacrylamide gel electrophoresis. Several proteins appear enriched in the supernatant of the LPS^*K.p*.^-treated sample (e.g., 12, 24, 37 and 60 kDa). The 24-kDa protein was identified as CadA (arrow). (**B**) Representative images of ETs elicited from parental AX4 and *cadA*^-^ mutant amoebae by LPS^*K.p*.^ exposure and stained with anti-CadA monoclonal antibodies (MLJ11, 1:10,000 dilution) followed by rabbit anti-Mouse AlexaFluor 488 antibodies (green). Fluorescent microscopy of ETs showing localization of CadA on the blue ET DNA scaffold that stains with DAPI. Bars, 20 µm.

### Bacterial binding to ETs

ETs were produced as above. Individual ETs were imaged by fluorescence microscopy. Bacteria bound was quantitated by creating a mask from the DAPI channel and the number of bacteria overlapping was counted. More than 30 individual ETs from three technical replicates were counted for each condition.

### ET-dependent killing by *D. discoideum* slug cells

AX4 and *cadA*^-^ slugs were developed as above and aliquoted into 24-well plates. To elicit ETs, LPS was added to a final concentration of 5 µg ml^-1^ and allowed to incubate for 15 minutes. *K. pneumoniae* was then added to a final amoeba:bacteria ratio of 100:1. Whole wells were removed and disaggregated at 30-minute intervals for two hours. Samples were incubated with the bacterium-permeable dye SYTO9 (Invitrogen) for 10 minutes before the bacteria were counted.

### Statistics

ET-dependent killing significance was calculated by Wilcoxon rank sum test.

## Results

We began to characterize ET-associated proteins in *D. discoideum* by purifying ETs elicited from Sentinel cells. After exposing the cells to lipopolysaccharide from the bacterium *Klebsiella pneumoniae* (LPS^*K.p*.^), we separated ETs from the cells using low-speed centrifugation (Methods). Our crude purification procedure results in a cell pellet and a supernatant fraction that is mainly ETs (11). After resolving the proteins in these fractions by gel electrophoresis, a number of proteins appeared to be enriched in the ET fraction produced by cells exposed to LPS^*K.p*.^ and the most prominent of those was a 24-kDa (Fig. 1A). We identified the protein by mass spectrometry as CadA, a *D. discoideum* developmental adhesion protein (18, 19). To determine whether or not CadA is actually present on ETs we used a monoclonal antibodies directed against CadA (MLJ11) to stain ETs that were freshly produced by Sentinel cells. ETs elicited from the wild-type strain (AX4) displayed uniform staining with the MLJ11 antibodies while ETs produced by the Sentinel cells of *cadA*^-^ mutants had no detectable signal (Fig. 1B). This indicated that our ET purification procedure correctly identified CadA as an ET-associated protein. In addition to its role in cell-cell adhesion during development, we have recently shown that CadA is also a lectin that is required for the establishment of a protective interface between vegetative amoebae and food bacteria (16). Current evidence suggests that the interface forms by CadA binding to the bacteria and agglutinating them, allowing the amoebae to feed on a matrix of bacteria without admixture of the two species (16). These findings suggested to us that CadA could be involved in bacterial interactions on ETs.

To test the potential role of CadA on ETs we first examined whether or not bacterial binding to ETs requires CadA. We used several species of bacteria expressing Green Fluorescent Protein (GFP) and incubated them with ETs elicited from wild-type (AX4) and *cadA*^-^ mutant Sentinel cells. We examined the bacteria localized to ETs using fluorescence microscopy and found significantly higher numbers of K. pneumoniae bacteria localized to wild-type ETs compared with *cadA* mutant ETs. We looked at dozens of ETs in several independent experiments carried out over months. Each wild-type ET that we examined had *K. pneumoniae* bacteria bound across most of its surface (Fig. 2A), whereas only occasional bacteria were observed bound to *cadA* mutant ETs (Fig. 2B). We quantified the bacterial binding by direct visual inspection (Fig. 2D). We observed similar pattern of localization of *E. coli* B/r bacteria to wild-type and *cadA* mutant ETs (Fig. 2D). This suggests that CadA is necessary for efficient binding of *K. pneumoniae* and *E. coli* B/r to ETs. To test this more directly, we added exogenous CadA to the binding assay during the elicitation of ETs from *cadA*^-^ mutant Sentinel cells. The added CadA restored the binding of *K. pneumoniae* and *E. coli* B/r to *cadA*^-^ mutant ETs to close to what we observed for wild-type ETs (Fig. 2C, D). The compromised binding of these two bacteria to *cadA*^-^ mutant ETs and the restoration of binding by exogenous CadA suggests that CadA acts as an ET adhesion protein for *Enterobacteriaceae*. We also tested the binding to ETs of two other species of bacteria that are not *Enterobacteriaceae*.

**Figure 2.**
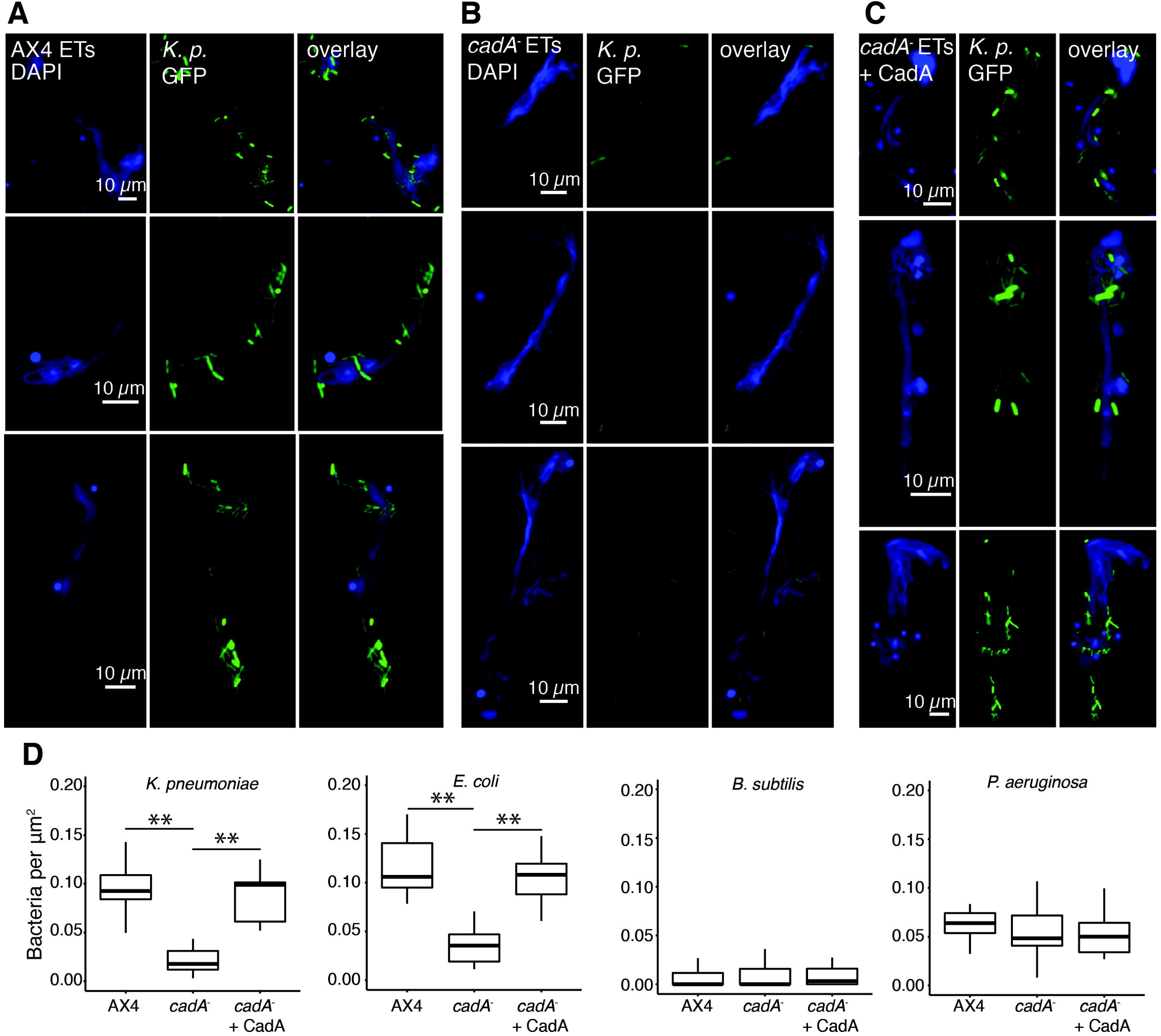
CadA requirement for localization of specific bacterial species to ETs. Wild-type (AX4) and *cadA* mutant disaggregated slug cells were incubated with *K. pneumoniae* GFP, *E. coli* B/r, *B. subtilis* GFP, or *P. aeruginosa* GFP on coverslips to elicit ETs. Note the co-localization of *K. pneumoniae* across the entire extent of AX4 ETs (**A**) but not on *cadA*^-^ ETs (**B**). (**C**) The addition of exogenous CadA restores localization of *K. pneumoniae* bacteria to *cadA*^-^ ETs. (**D**) Box plots showing CadA dependence of *K. pneumoniae* and *E. coli* B/r binding to ETs, *B. subtilis* not binding ETs, and *P. aeruginosa* binding ETs independent of CadA. Data represent three independent experiments (** p<0.005).

Interestingly, we found that *Bacillus subtilis* did not localize to wild-type or *cadA*^-^ mutant ETs, while *Pseudomonas aeruginosa* localized equally well to both the wild-type and *cadA*^-^ mutant ETs (Fig. 2D). Additionally, the localization of *B. subtilis* or *P. aeruginosa* to ETs did not increase or decrease when we added CadA protein (Fig. 2D). These results suggest that CadA promotes the adhesion to ETs of particular species of bacteria, possibly only *Enterobacteriaceae*.

Next, we examined the physiological significance of CadA-stimulated ET localization by measuring ET-dependent and CadA-dependent killing of bacteria. We incubated preparations of slug cells with *K. pneumoniae* and quantified bacterial killing by *BacLight*™ LIVE/DEAD staining and fluorescence microscopy. We have previously demonstrated that the number of live bacteria estimated in this way correlates with the number viable bacteria, as determined by their ability to form colonies on agar growth plates (19). We observed efficient bacterial killing (>85%) by AX4 slug cells, whereas *cadA*^-^ mutant slug cells killed ∼50% of the bacteria (Fig. 3A).

**Figure 3.**
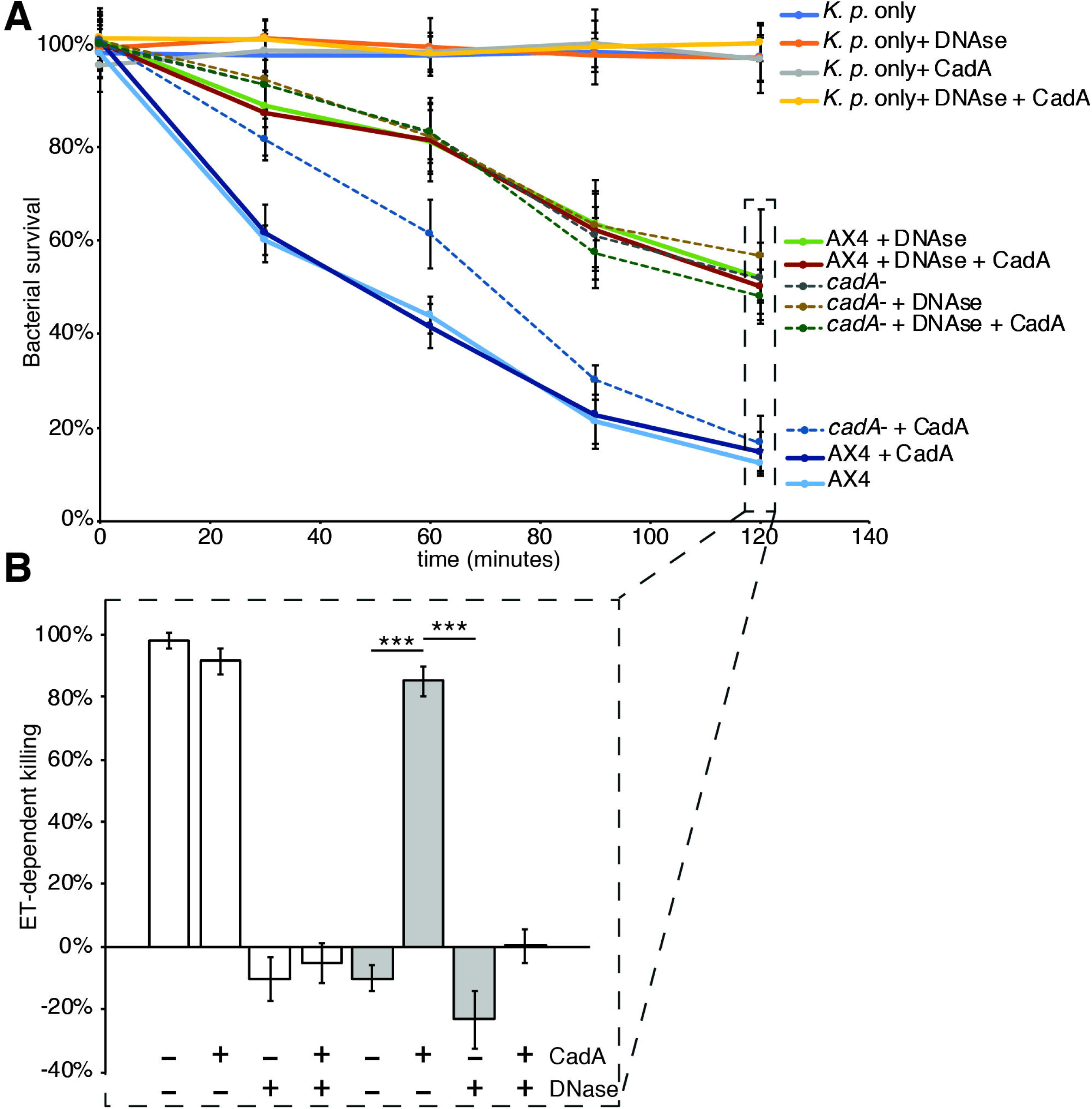
CadA protein restores bacterial killing by *cadA^-^* ETs. (**A**) *K. pneumoniae* bacteria were exposed to AX4 (solid lines) or *cadA*^-^ (dashed lines) slug cells and *BacLight*™ LIVE/DEAD staining was used to measure viable bacteria over time. Data was normalized to *K. pneumoniae* only samples (top, solid blue line) and are the result of three technical replicates of three biological replicates. (**B**) ET-dependent killing at 120 min as determined by subtracting the average bacterial survival in the AX4 + DNAse samples then normalizing to AX4 with no additions. *** p<0.0001, Wilcoxon rank sum test.

To assess ET-dependent killing, we carried out the assay in the presence of DNase which rapidly destroys the ET structure and presumably dilutes the ET-associated antimicrobials into the assay medium (6, 11). The addition of DNase reduced bacterial killing by AX4 cells to the level we observed in *cadA*^-^ mutant cells, but had little effect on *cadA*^-^ mutant bacterial killing (Fig. 3A). Therefore, we defined ET-dependent killing as the difference between the bacterial killing we observed by slug cells before and after the addition of DNase (Fig. 3B). This is consistent with our observations of the absence of ET-dependent killing of bacteria by *cadA*^-^ mutant ETs determined by microscopic examination. Also, our observation of ET-independent killing suggests that the genetic ablation of *cadA* does not affect other antimicrobial functions of slug cells in our assay. To test if exogenous CadA could rescue ET-dependent killing by *cadA* mutant slug cells, we added CadA to our assay during ET elicitation. Supplementing *cadA* mutant slug cells with purified CadA increased bacterial killing to wild-type levels and it did not affect killing by wild-type cells (Fig. 3). To confirm that the effect of CadA was caused by an increase in ET-dependent killing we added DNase along with CadA during ET elicitation. Added DNase brought the CadA-enhanced bacterial killing by *cadA*^-^ mutant cells back to the original level we observed for *cadA*^-^ mutant cells alone, indicating CadA affected ET-dependent killing in our assay. The control assays, without added slug cells, showed that we could recover 100% of the bacteria when DNase, CadA, or both, were added to the assay, demonstrating that the loss of bacteria was not due to killing by DNase or because they became unrecoverable due to CadA agglutination (Fig. 3A).

## Discussion

Here we showed that a bacterial agglutinin CadA acts as a bacterial adhesion protein on *D. discoideum* ETs. CadA is necessary and sufficient for *K. pneumoniae* and *E. coli* B/r to bind to ETs, and for ET-dependent killing of *K. pneumoniae*. Other bacterial species that we tested are agnostic to the presence of CadA on ETs suggesting additional factors may exist handle other bacterial species. Indeed, two CadA orthologs, Cad2 and Cad3, are expressed in the slug (20, 21), with 30% and 73% amino acid identity respectively, and may represent additional ET-associated bacterial adhesion proteins. Bacteria also express adhesins (22, 23), pili (24), and secretion systems (25) that target host surface proteins or glycans (26). Perhaps host targets of those bacterial proteins are packaged onto ETs to provide additional mediators of bacterial adhesion and killing on *D. discoideum* ETs that is independent of Cad proteins.

Structural studies revealed that CadA has a C-terminal Ig-like domain and an N-terminal gamma-crystallin-like fold with three Ca^2+^ binding sites (27). Cell-cell adhesion by CadA is accomplished through homotypic head to head (N-to-N) binding, with the Ig-like domains binding to cell surface protein proteins on adjacent amoebae. Importantly, incubating CadA with EGTA inhibits amoebal adhesion and bacterial agglutination but does not interfere with CadA binding to bacteria (16). This suggests that the gamma-crystallin-like fold is required for dimerization of CadA and the Ig-like fold contains the carbohydrate (lectin) binding activity. From this information we would speculate that the CadA gamma-crystallin-like domain associates with the ET (or an ET-associated protein) while the Ig-like domain binds to the bacteria, likely a surface glycan. CadA is present within the cytoplasm of the amoeba and is secreted through the contractile vacuole (28). This is accomplished through association with a contractile vacuole membrane protein that inverts, translocating CadA into the lumen of the contractile vacuole. Association of CadA with ETs may occur by either by extracellular self-assembly, given the high level of CadA expression in slugs (20, 21), or by its packaging with mtDNA during ET maturation within the cell.

The survival benefit of ETs is thought to lie in their ability to bind and kill bacteria efficiently by delivering a high local concentration of antimicrobials at the correct time and place in the body. Consistent with this idea, hydrolyzing the ET DNA scaffold reduces bacterial killing suggesting that binding to ETs is a critical aspect of innate immune function. While many studies have correctly focused on the antimicrobial components of ETs, bacterial adherence to ETs has been overlooked. Localized concentrations of antimicrobials provide little benefit if bacteria can avoid the location. Since carbohydrate binding lectins are used by the mammalian innate immune system to detect and kill pathogens in other contexts (29, 30), it is worth considering that they are also needed for bacterial adherence to mammalian ETs.

## Acknowledgement

We thank Olga Zhuchenko for her observations of the enrichment of CadA in ET preparations shown in Figure 1A, and William F. Loomis and Gad Shaulsky for providing insights during the course of this work, and Danny Fuller and WFL for the gift of MLJ11 antibodies.

## Author contributions

TF and AK conceived and designed the experiments, TF directed and performed all of the experiments with assistance from CD. TF and AK wrote the manuscript with scientific and editorial input from CD.

